# Validation of RT-qPCR approaches to monitor *Pseudomonas syringae* gene expression during infection and exposure to pattern-triggered immunity

**DOI:** 10.1101/183269

**Authors:** Amy Smith, Amelia H. Lovelace, Brian H. Kvitko

## Abstract

*Pseudomonas syringae* pv. tomato DC3000 (DC3000) is an important model plant pathogen, with a fully annotated genome and multiple compatible plant hosts. Very few studies have examined the regulation of DC3000 gene expression *in vivo*. We developed a RT-qPCR assay to monitor transcriptional changes in DC3000 inoculated into *Arabidopsis thaliana* leaves during disease and exposure to pattern-triggered immunity (PTI). In our approach, bacterial RNA concentrations in total tissue RNA are standardized using *P.syringae*-specific16S ribosomal RNA primers. We validated multiple stable reference genes for normalization in calculating the relative expression of genes of interest. We used empirically derived rates of amplification efficiency to calculate relative expression of key marker genes for virulence-associated regulation. We demonstrated that exposure to PTI alters DC3000 expression of Type III secretion system, coronatine synthesis genes and flagellar marker genes.

## Introduction

*Pseudomonas syringae* pv. tomato DC3000 (DC3000) is a commonly used and powerful model bacterial pathogen of plants. DC3000 is both highly genetically tractable and benefits from a complete genome sequence that has undergone continuous efforts to update and refine its annotation (Buell, Joardar et al. 2003, Lindeberg, Biehl et al. 2009, Filiatrault, Stodghill et al. 2011). The capacity of DC3000 to infect both the model plant hosts *Arabidopsis thaliana, Nicotiana benthamiana*, and the horticultural crop tomato *(Solanum lycopersicum)*, have made it a preferred pathogen for analysis of plant biotic stress, plant microbe interactions, and the study of the plant immune system (Zeng, Brutus et al. 2011, Bombarely, Rosli et al. 2012, Xin and He 2013). Pattern-triggered immunity (PTI), one of two major tiers of the plant immune system, confers broad defense against pathogens of multiple classes, typically in the absence of a programmed-cell-death hypersensitive response (Jones and Dangl 2006, Nishimura and Dangl 2010, Zipfel and Robatzek 2010, Newman, Sundelin et al. 2013). PTI is activated by the binding of Pathogen/Microbe Associated Molecular Patterns (P/MAMPs) such as microbial cell wall components or epitopes of conserved microbial proteins by cognate surface displayed Pattern Recognition Receptors (PRRs) (Jones and Dangl 2006). In Arabidopsis, the 22 amino acid peptide epitope of bacterial flagellin, Flg22, is recognized by the PRR FLS2 leading to the activation of PTI in the plant host (Zipfel, Robatzek et al. 2004). In Arabidopsis, pre-induced PTI restricts DC3000 proliferation and the delivery of protein effectors by the DC3000 Type III Secretion System (T3SS) (Crabill, Joe et al. 2010, Wei, Chakravarthy et al. 2013). The mechanism(s) by which PTI restricts T3SS effector delivery are unclear. A potentially fruitful approach that could be used to uncover the mechanisms of PTI would be to monitor DC3000 transcriptomic responses *in vivo* during PTI exposure.

A handful of transcriptomic approaches have been used to examine the global gene expression patterns of *P. syringae in vivo*. Genetic screening using *In Vivo* Expression Technology (IVET) revealed differential expression of many DC3000 genes now known to be virulence-associated, as well as many other plant-induced genes whose roles in plants have yet to be determined (Boch, Joardar et al. 2002). Similarly, microarray analysis has been used to identify *P. syringae* pv. *syringae* B728a genes differentially regulated during epiphytic or endophytic colonization of bean (Yu, Lund et al. 2013).

Quantitative, reverse-transcription polymerase chain reaction (RT-qPCR) is a well-established and powerful tool for small-scale transcriptomic studies. As a complement to RNASeq, RT-qPCR allows researchers to investigate transcriptional changes between experimental conditions in a more focused manner. RT-qPCR has been used routinely to monitor *P. syringae* gene expression *in vitro* in hrp-inducing media (Wei, Plovanich-Jones et al. 2000, Tegli, Gori et al. 2011, Vargas, Farias et al. 2013, Lee, Ryu et al. 2015) but only a few studies have examined gene expression *in vivo* (Mascia, Santovito et al. 2010, Tegli, Gori et al. 2011, Scholtz and Visser 2013, Petriccione, Mastrobuoni et al. 2015). The study of bacterial transcriptomics within hosts is not straightforward due to the low and variable amounts of bacterial RNA present across stages of infection amidst host and endogenous microbiota RNA. These wide variations in bacterial RNA concentration can make it difficult to standardize all samples to a single bacterial RNA concentration across diverse *in planta* treatments. In addition, test-condition-appropriate reference genes should be validated as stable for normalizing expression of genes of interest. Without appropriate sample standardization and validated expression normalization, relative expression calculation for target genes of interest (GOIs) can be skewed.

We sought to address these problems by validating a RT-qPCR method to measure DC3000 gene expression directly in inoculated Arabidopsis leaves, following the Minimum Information for Publication of Quantitative Real-Time PCR Experiments (MIQE) guidelines (Bustin, Benes et al. 2009). The MIQE guidelines were published in 2009 to establish a rubric regarding all aspects of RT-qPCR, from sample preparation to data storage. To develop our approach, we generated *in planta* samples of different treatment conditions, including early (5 h) and middle (24 h) infections by DC3000 in Arabidopsis Col-0 leaves either pre-treated 20 h prior with Flg22 to activate PTI, or mock treated to allow natural disease progression. We developed *P. syringae-specific* primers for the 16S ribosomal small subunit RNA (16S rRNA) to measure the concentrations of bacterial RNAs within mixed RNA samples. We tested the transcriptional stability of six published DC3000 reference genes (16S rRNA, *gap-1, gyrA, hemD, recA, and rpoD)* (Ferreira, Myers et al. 2006, Records and Gross 2010, Tegli, Gori et al. 2011, González-Lamothe, El Oirdi et al. 2012, Freeman, Chen et al. 2013, Vargas, Farias et al. 2013, Chatnaparat, Li et al. 2015, Ishiga and Ichinose 2015, Lee, Ryu et al. 2015) as well as three new reference gene candidates selected based on a preliminary *in situ* RNAseq experiment, ultimately validating four for use in our approach *(recA, rpoD, leuD, and oprF)*. Using our protocol for RT-qPCR with multiple validated RGs, we examined the relative expression of three GOIs *(hrpA, cfl* and *fliC)* relevant to DC3000 host interactions of DC3000 *in planta*, and from *in vitro* media.

## Results

### Quantification of bacterial RNA from inoculated tissue total RNA

For accurate calculation of relative gene expression by RT-qPCR, it is important to standardize to a single concentration of bacterial RNA across samples. During the course of infection, DC3000 colony counts within leaf tissue may range by three or more orders of magnitude. One approach that could be used to estimate bacterial RNA in a total tissue RNA sample would be to determine viable bacterial counts by dilution plating and standardize input RNA based on those bacterial counts. Unfortunatley, viable count sampling must be conducted from a secondary tissue sample as the time and processing needed for dilution plating would result in alterations to the bacterial transcriptome. Conversely, RNA stabilizing techniques would interfere with accurate measurement of viable colony counts. Thus colony counts must be determined from a second sample collected from the same inoculated tissue used for RNA extraction. We wanted to validate a bacterial RNA quantification method directly linked to the RNA sample to be RT-qPCR-analyzed, rather than a method that requires inference from a second tissue sample. By quantifying directly from the analytical sample, any variations between samples in RNA extraction and cDNA conversion are directly incorporated into the RNA quantification measurement. We tested whether DC3000 16S rRNA concentration could be used as an in-sample proxy for bacterial concentration within inoculated tissue, using the rationale that ribosomal RNA comprises the major component of total bacterial RNA (Bremer and Dennis 2008) and thus could serve as a reliable measurement of bacterial RNA concentration as a whole. To avoid amplification of endogenous microbiome or chloroplast 16S rRNA, specialized *P. syringae* 16S primers were designed to prime within *P syringae* 16S variable regions 1 and 2 to improve specificity. In Pseudomonads, variable region 1 is hypervariable, allowing for good species specificity in primer design (Bodilis, Nsigue-Meilo et al. 2012).

To test whether bacterial 16S rRNA levels detected by RT-qPCR could be used as a proxy for measuring bacterial concentration *in planta*, plants were inoculated with DC3000 suspensions at five concentrations and then sampled 5 hours post inoculation (hpi). To determine bacterial concentration and to extract total RNA, bookmatched leaf discs punched from left and right sides of individual leaves were carefully distributed for either dilution plating or RNA extraction. For each sample, we paired the number of viable bacteria per mg leaf tissue (CFU/mgleaf tissue) as determined by dilution plating with the concentration of bacterial 16S rRNA (ng 16S rRNA/mg leaf tissue) determined by RT-qPCR using *P. syringae-specific* 16S rRNA primers. 16S rRNA concentration was determined using a standard curve generated with a synthetic double-stranded DNA gene block (gBlock) of the target 16S rRNA amplicon. Pairing these two metrics (CFU/mg leaf tissue and ng 16S rRNA/mg leaf tissue, both log10-transformed) against one another for each sample yields a positive linear correlation of CFU per ng bacterial 16S rRNA (Fig 1A).

**Figure 1.**
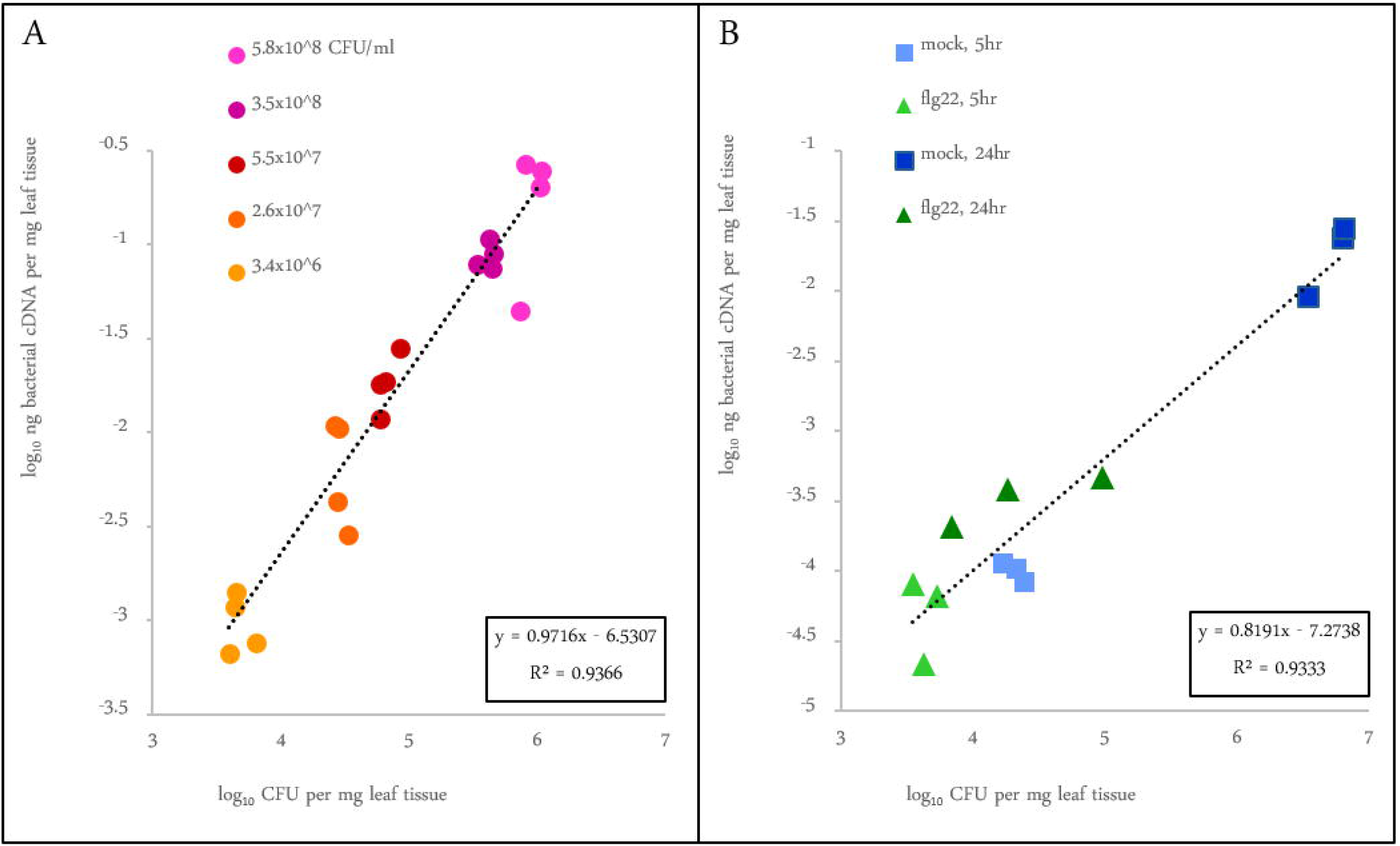
Correlation of bacteria counts to quantities detected by RT-qPCR. A) Decreasing inoculum concentration results in equivalent detection by RT-qPCR. *Arabidopsis* leaves were syringe-infiltrated with the indicated concentrations of inoculum, and sampled at 5 hpi. Thirty-two 0.125 cm2 discs were punched from infected leaves and split for both testing methods. Bacterial CFU per mg leaf weight was determined by dilution plating; in the partnered samples, ng 16S rRNA/mg leaf tissue was calculated by RT-qPCR from a standard curve. Pairing the log_10_ of each of these measurements yields a strong positive linear correlation with a slope of 0.972 and a coefficient of determination (R^2^) of 0.9366. Markers on the graph represent individual biological replicates (n=4). **B) 16S rRNA can be used as an effective proxy for bacterial load in PTI-preactivated tissue and at different stages of infection**. Plants pre-treated 20 hpi with 1 μM Flg22 in 0.1% DMSO and 0.25 mM MgCl_2_ were protected from *P. syringae* infection through pre-activation of PAMP-triggered immunity. Mock-treated plants received 0.1% DMSO in 0.25mM MgCl_2_ only and remained susceptible to infection. Plants were infected with 1.9x10^8^ CFU/ml inoculum and sampled at 5 and 24 hpi. Bacterial CFU per mg leaf weight was paired with ng 16S rRNA per mg leaf weight as detected by RT-qPCR. The log_10_ of each of these measurements still yields a positive linear correlation, with a slope of 0.819 and a coefficient of determination (R^2^) of 0.9333. Markers on the graph represent individual biological replicates (n=3).

It is possible that ribosome content per bacterium could vary during *in planta* treatments or during exposure to PTI. This would prevent us from using 16S rRNA concentration as an accurate proxy for bacterial concentration. We conducted an experiment to test whether a linear relationship between bacterial count and 16S rRNA is retained in cells exposed to *in planta* conditions that either restrict or permit bacterial growth. In this experiment, cell numbers and 16S RNA levels were determined for DC3000 in mock or Flg22-pretreated leaves were inoculated with one inoculum concentration and samples were collected at 5 and 24 hpi. Bacterial CFU/mg counts were again paired with RT-qPCR-determined 16S rRNA quantities (Fig 1B). This demonstrated a similar relationship between bacterial counts and 16S RNA per mg leaf tissue. As expected, the concentration of bacteria in Flg22-pretreated plants did not increase over the course of 24 h, whereas during infection of mock-treated plants the concentration of bacteria increased three logs over the same period of time (Fig 1B). Even though there was a difference in growth, these two populations displayed similar per cell rRNA ratios. These results support our approach of using RT-qPCR-calculated 16S rRNA concentration as a direct *in situ* estimate of bacterial RNA concentrations in mixed samples to standardize bacterial RNA input in later steps.

### Reference gene selection

Reference genes (RG) are necessary to normalize measurements of transcript abundance between samples or experimental treatments. MIQE guidelines call for the use of multiple RGs in calculating transcriptional changes. The ideal RG set would remain transcriptionally stable under different experimental conditions. MIQE quidlines also recommend pairing GOIs with RGs of similar expression values, thus having validated RGs covering a wide range of expression values allows for flexibility of pairing with high- or low-expressed GOIs. We identified nine RG genes to test based on historical utility or new global expression data that suggest they might be ideal candidates for use as reference genes in these types of studies. Previous studies of *P. syringae* have used *gyrA, gap-1, hemD, recA* and *rpoD* and *16S rRNA* as individual RGs (Ferreira, Myers et al. 2006, Records and Gross 2010, González-Lamothe, El Oirdi et al. 2012, Vargas, Farias et al. 2013, Ishiga and Ichinose 2015, Lee, Ryu et al. 2015) (Tegli, Gori et al. 2011, Freeman, Chen et al. 2013, Chatnaparat, Li et al. 2015). *gap-1* and *gyrA* showed equal expression under *in vitro* experimental conditions as measured by microarray (Ferreira, Myers et al. 2006). We used previously-published qPCR primers for *hemD* (Freeman, Chen et al. 2013) and *rpoD* (González-Lamothe, El Oirdi et al. 2012), and designed new primers to target *gyrA, gap-1* and *recA*. The remaining four RG candidates, *cynT, lcs-1, leuD* and *oprF* were observed as highly expressed and stable in Flg22-pretreated and mock-treated infected plant tissues from a preliminary RNAseq analysis (manuscript in preparation). We designed new primers for these four candidate RGs as well (Table 1).

**Table 1.**
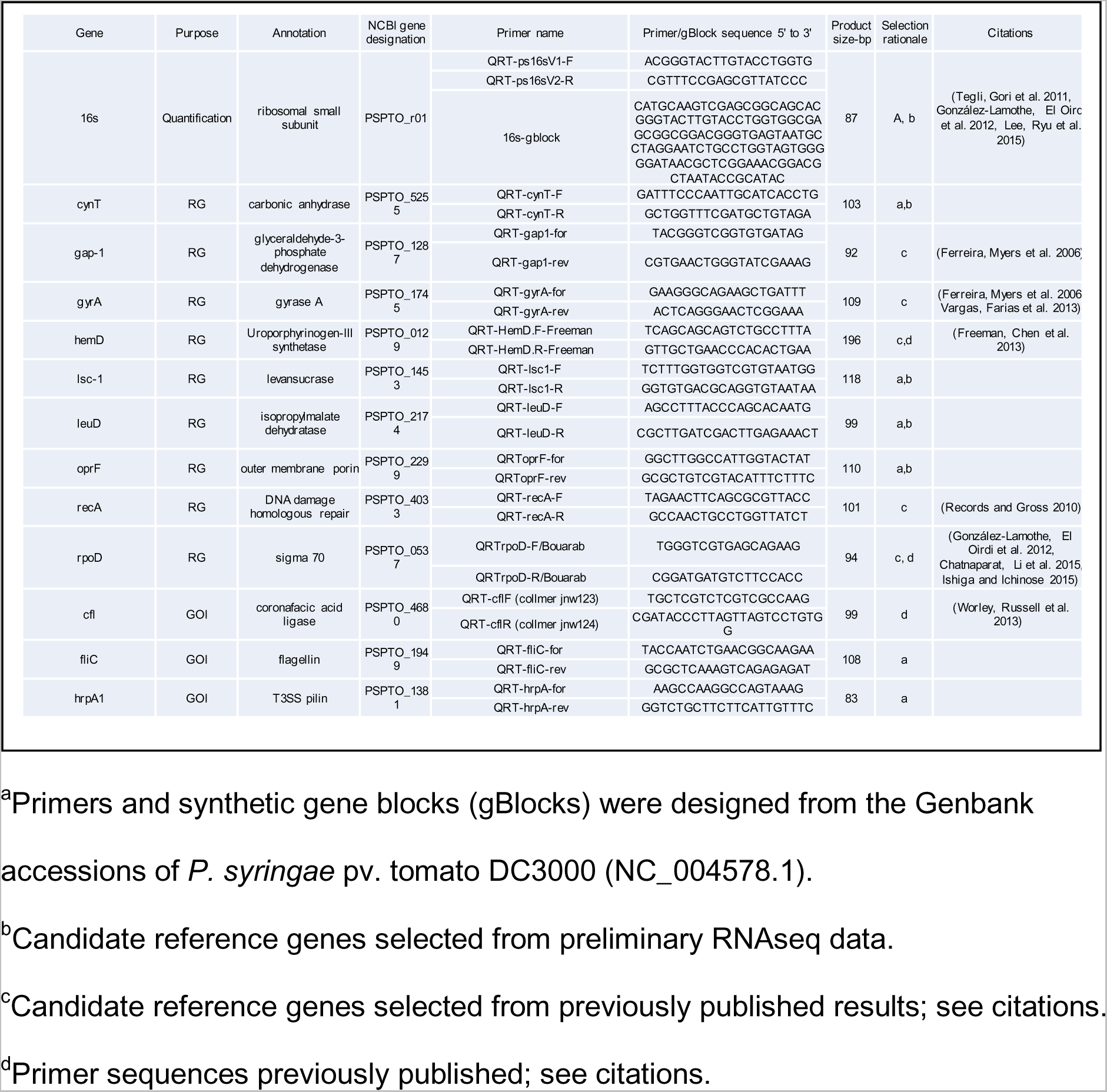
Target genes and corresponding sequences for primers and gBlocks.

### Reference gene validation and efficiency estimates

The efficiency of PCR is the percentage of perfect doubling of amplicons from cycle to cycle. It can be measured in one of two ways: either from the slope of a curve generated by pairing known quantities (usually serially-diluted) of target cDNA to their resulting quantification cycle (Cq) values of the PCR, or by directly measuring the change in fluorescence per cycle from raw PCR data. The latter is a more direct method, and is derived empirically from each sample during RT-qPCR by inputting raw fluorescence data into the software LinRegPCR (LRP), available from the Heart Failure Research Center. We compared the PCR efficiency as estimated by the serial-dilution method (Eest) to the efficiency derived empirically in LRP (Eemp) for all primer sets used in this study. Eest was generally 5-10% higher than its partner Eemp, and often estimated to be greater than 100% (Table 2; Supplementary data S1, “RG pre-tests”).

**Table 2.**
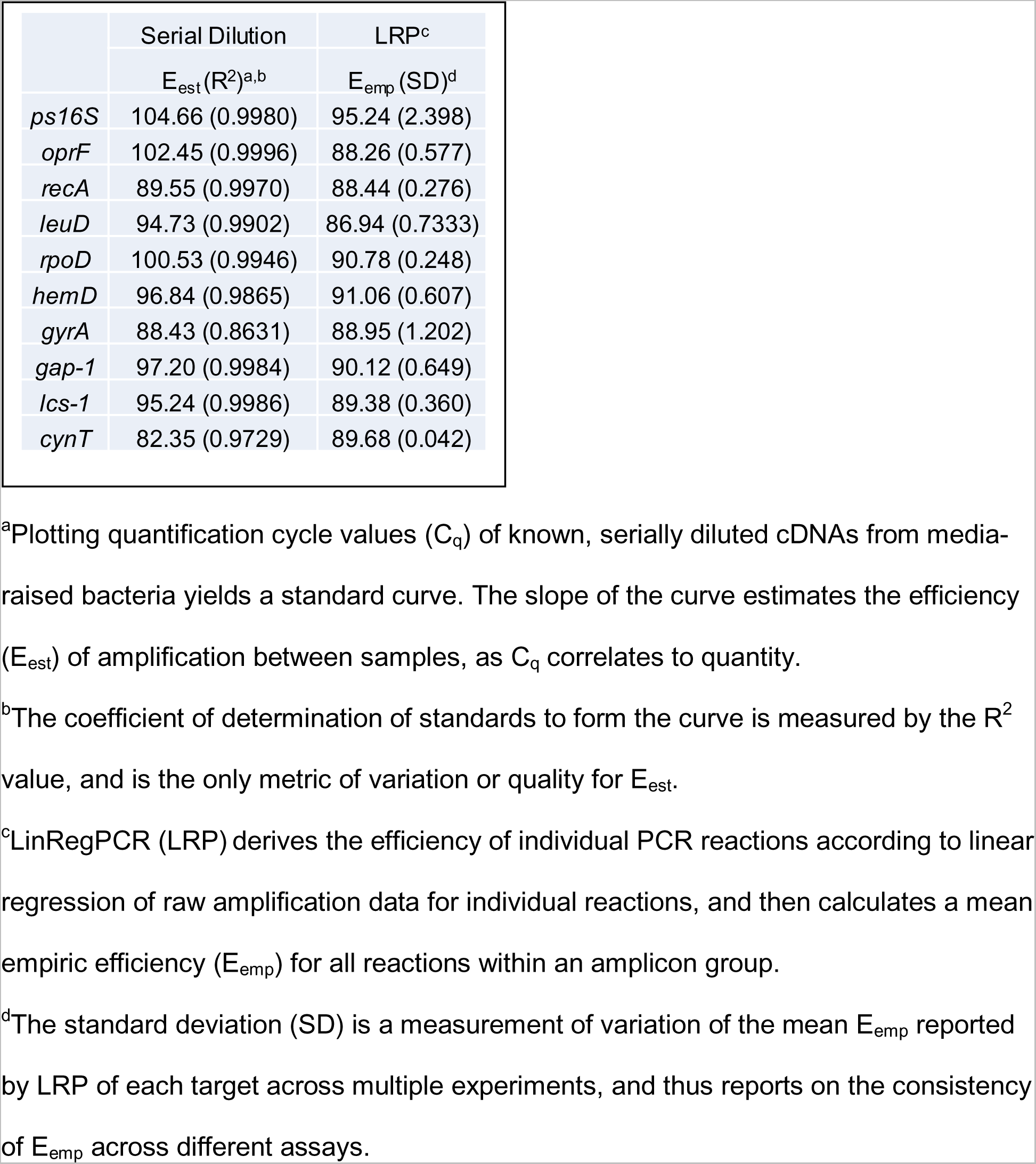
Estimated Efficiency and Empiric Efficiency of RT-qPCR primer sets used in this study.

|  | Serial Dilution | LRP<sup>c</sup> |
| --- | --- | --- |
|  | E<sub>est</sub> (R<sup>2</sup>)<sup>a,b</sup> | E<sub>emp</sub> (SD)<sup>d</sup> |
| *ps16S* | 104.66 (0.9980) | 95.24 (2.398) |
| *oprF* | 102.45 (0.9996) | 88.26 (0.577) |
| *recA* | 89.55 (0.9970) | 88.44 (0.276) |
| *leuD* | 94.73 (0.9902) | 86.94 (0.7333) |
| *rpoD* | 100.53 (0.9946) | 90.78 (0.248) |
| *hemD* | 96.84 (0.9865) | 91.06 (0.607) |
| *gyrA* | 88.43 (0.8631) | 88.95 (1.202) |
| *gap-1* | 97.20 (0.9984) | 90.12 (0.649) |
| *lcs-1* | 95.24 (0.9986) | 89.38 (0.360) |
| *cynT* | 82.35 (0.9729) | 89.68 (0.042) |

<sup>a</sup>Plotting quantification cycle values (C<sub>q</sub>) of known, serially diluted cDNAs from media-raised bacteria yields a standard curve. The slope of the curve estimates the efficiency (E<sub>est</sub>) of amplification between samples, as C<sub>q</sub> correlates to quantity.
<sup>b</sup>The coefficient of determination of standards to form the curve is measured by the R<sup>2</sup> value, and is the only metric of variation or quality for E<sub>est</sub>.
<sup>c</sup>LinRegPCR (LRP) derives the efficiency of individual PCR reactions according to linear regression of raw amplification data for individual reactions, and then calculates a mean empiric efficiency (E<sub>emp</sub>) for all reactions within an amplicon group.
<sup>d</sup>The standard deviation (SD) is a measurement of variation of the mean E<sub>emp</sub> reported by LRP of each target across multiple experiments, and thus reports on the consistency of E<sub>emp</sub> across different assays.

To measure the stability of the RGs, the differently treated *in planta* samples from a preliminary experiment (similar to that in Fig 1B) and an *in vitro* sample were standardized for bacterial RNA using 16S rRNA. These samples were RT-qPCR-amplified with primers targeting all nine candidate RGs; *P. syringae-specific* 16S rRNA primers were also used to confirm equal loading. The raw amplification data was input to LRP, and the C_q_ outputs were measured for variability by RefFinder, available from the Cotton EST Database. Candidate RGs were classified by C_q_ range (high expressed: 15 – 25; mid expressed: 20 – 30; low expressed: 25 – 35) using the observed Cqs from preliminary data (Supplementary data S1, “RG pretests”).The candidate RGs were ranked from most stable (least variability among standardized samples) to least stable (most variability among standardized samples). Of the nine candidate RGs, *oprF, leuD, recA*, and *rpoD* were found to be the most stable; these four genes had valid C_q_ ranges from 14-30 as assessed by R^2^ values of data plotted from E_est_ serial dilutions (Figure 2, Supplementary data S1.)

**Figure 2.**
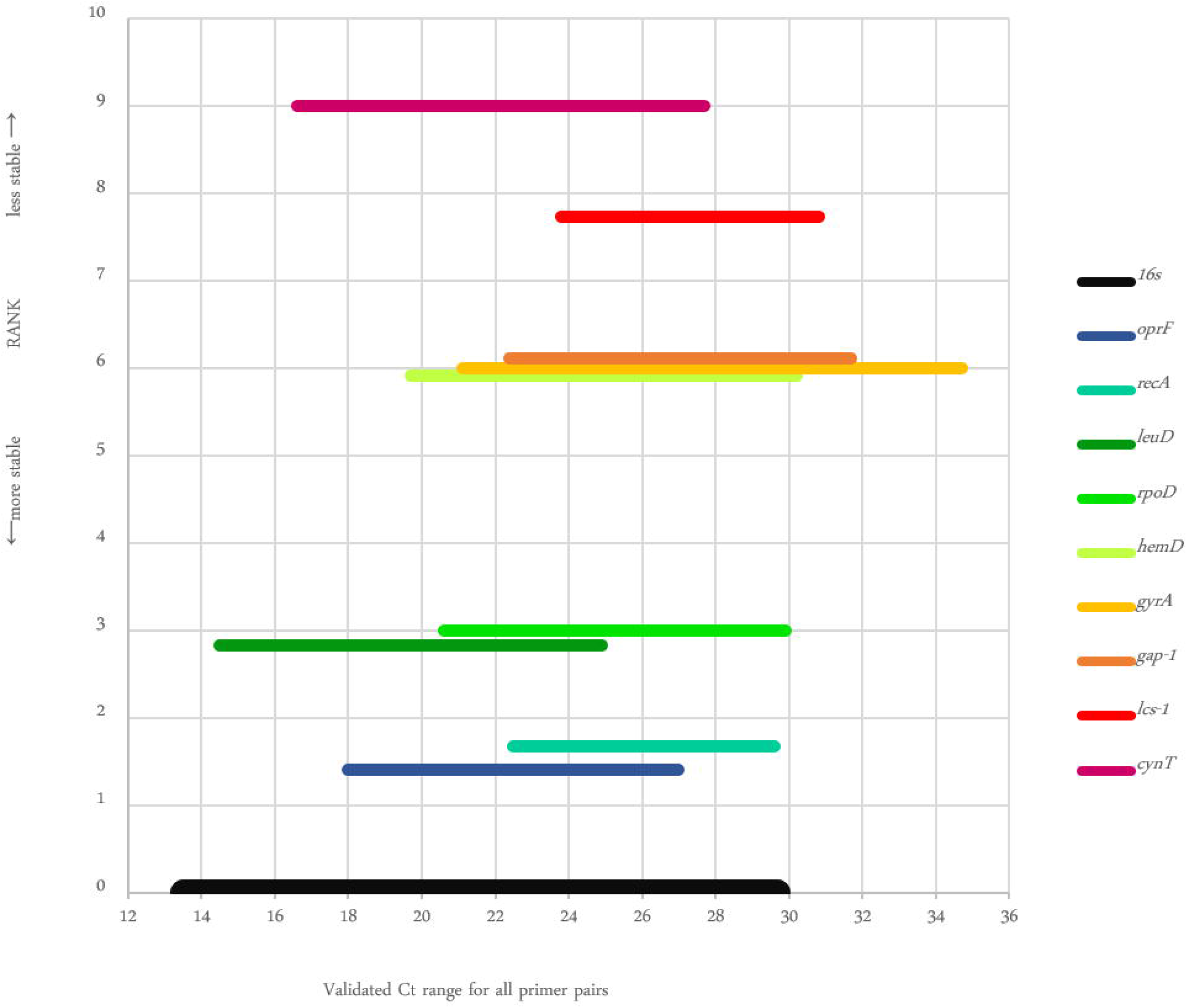
Comprehensive Gene Stability rank and Cq ranges of candidate reference genes as determined using RefFinder. Y *axis*: larger values indicate decreasing stability and poor scores, ranked by the geometric mean of scores according to similarity of Cq values for templates of equal concentration. *X axis*: Range of functional Cq for each reference gene, as determined by a preliminary run of RT-qPCR against varying concentrations of cDNA from DC3000 grown on KB media. 16S rRNA is represented here to show its Cq range but was not included in stability calculations.

### Using multiple reference genes

It is increasingly recommended that more than one RG be used when quantifying relative expression as there is no “one true” stable reference gene that is compatible with every given GOI or experimental condition (Vandesompele, De Preter et al. 2002, Bustin, Benes et al. 2009, Kozera and Rapacz 2013). To determine whether the use of single, paired, or multiple RGs would significantly change calculations of the relative expression of a gene of interest, we calculated the Normalized Relative Quantity (NRQ) using different combinations of RGs. Three biological replicates each of the 24 hpi, mock *in planta* samples and the *in vitro* bacteria samples were standardized to a single 16S rRNA quantity (0.6pg) and RT-qPCR-amplified against the four validated RGs and 16S rRNA. From the resultant E_emp_ and C_q_ values derived by LRP, we calculated NRQ values using different numbers of RGs (five single RGs, four pairs of RGs, and two combinations of all RGs with and without 16S rRNA), and analyzed for significant differences by ANOVA (p<0.05). We found that except for 16S rRNA, any single RG or any combination of two or more of our four validated RGs gave NRQs that were not significantly different from one another. (Supplementary data S1, “RG pairs”). Based on this, we used *oprF and recA* together when testing for NRQ, because they represent normal and high C_q_ ranges, for pairing with high- and mid-expressed genes of interest, and because they satisfy the MIQE guideline to use more than one RG.

### Relative expression of key virulence-regulated genes during infection and PTI exposure

To test our approach, we monitored the expression of three key virulence-associated genes comparing expression *in vitro* from KB media to the *in planta* conditions of PTI or infection. We selected *hrpA*, (T3SS *hrp* pilin), *cfl* (coronafacic acid ligase), and *fliC* (flagellin) as marker genes of T3SS-associated genes, the jasmonic-acid-mimicking phytotoxin coronatine and the flagella, respectively. Total RNA from inoculated tissue was isolated after 5 hpi and 24 hpi in Arabidopsis plants pre-treated with Flg22 to pre-induce PTI or with a mock treatment for infection of naïve plants. RNA was also extracted from a sample of the bacterial inoculum prepared from KB media. Six experimental replicates of the *in planta* and *in vitro* DC3000 samples were standardized for equal loading of bacterial RNA based on 16S rRNA. We tested the three target GOIs against the two RGs, (*oprF, recA*) and used the *in vitro* inoculum sample as a calibrator of relative expression.

PTI exposure affected the regulation of each GOI in ways that suggest dynamic regulatory responses to PTI exposure and infection. First, *hrpA* was induced more strongly during PTI exposure than during infection at 24 hpi (Fig 3A), even thought it was similarly expressed both during infection and PTI exposure at 5 hpi (Fig 3A). Conversely, *cfl* was differentially induced during infection compared to PTI exposure at 5 hpi, but at 24 hpi *cfl* expression was similarly low in both treatments (Fig 3B). Relative expression of *fliC* was down regulated at both 5 and 24 hpi in both treatments, but significantly more repressed during infection than during PTI exposure (Fig 3C).

**Figure 3.**
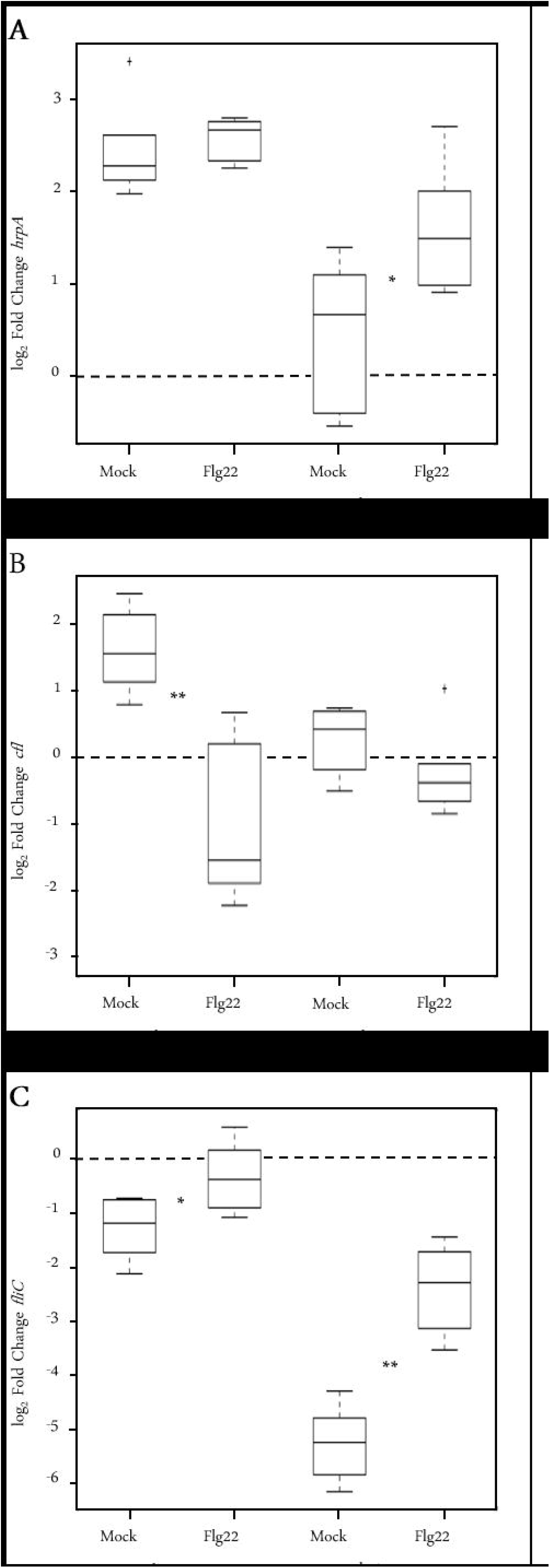
Relative Expression of three Genes of Interest. Expression of three genes of interest (GOI) of *in planta* samples relative to expression of DC3000 grown *in vitro* in KB media, normalized against the expression of two reference genes, *oprF* and *recA*. Each box plot is formed from six biological replicates, except Flg22, 5hr, for which n=5. T-tests (two-tailed, assuming equal variances) were conducted between mock and Flg22 samples for both time points. The dashed line indicates zero-fold change compared to *in vitro* DC3000. Single asterisks indicate significant differences with p<0.05; double asterisks indicate p<0.01. **A) T3SS pilin** At 5 hpi, expression of *hrpA* is up in both mock- and Flg22-pretreated bacteria, relative to media-raised bacteria, but there is no significant difference between treatments. By 24 hpi, expression of *hrpA* has reduced in both samples, more so in the mock-treated sample. **B) Coronafacic acid ligase** When compared to media raised bacteria, expression of *cfl* at 5 hpi is induced in mock-treated plants, but is repressed in bacteria exposed to Flg22-pretreated plants. No significant difference exists between mock and Flg22-pretreated leaves at 24 hpi. **C) Flagellin** *fliC* is negatively expressed in all *in planta* treatments, compared to media-raised bacteria. For both time points, *fliC* is significantly repressed in Flg22-pretreated samples compared to mock-treated plants.

### Comparison of methods to calculate NRQ

Early publications calculated relative expression using the 2^-ΔΔCt^ method described by Livak (Livak and Schmittgen 2001). In this calculation, “2” refers simultaneously both to the doubling of amplicons in each PCR cycle, as well as the efficiency, with 2 being “perfect doubling”. Livak’s equation assumes all PCRs to have perfect efficiency, which is rarely true. Pfaffl amended this equation to allow users to input unique efficiencies for both the GOI and the reference gene (Pfaffl 2001); this yields a more precise and usually more conservative measurement of relative expression. An additional benefit to this approach is that it allows meaningful comparisons of relative expression between different genes, which cannot be done if unless variations in efficiency are accounted for. One challenge, however, with the Pfaffl equation is that it does not allow users to utilize more than one RG. A modification (Vandesompele, De Preter et al. 2002) to the Pfaffl equation – using the geometric mean of the normalizing factors of multiple RGs – resolved this issue. Using the above data, we compared these two methods to calculate relative expression. NRQ calculated via the modified Pfaffl equation was more conservative and less variable compared to the calculation using Livak equation (Supplementary data S1, “ddCt vs Pfaffl”). While the difference in both measurements is slight, we chose to use the modified Pfaffl equation with LRP-transformed data, as it is a better, more precise approach toward presenting RT-qPCR data.

## Discussion

Our objective with this study was to validate a RT-qPCR approach to quantify relative expression of DC3000 transcripts *in planta* during infection and PTI-exposure. To achieve this, we designed *P. syringae-specific* 16S rRNA primers for RT-qPCR analysis on the hypothesis that ribosomal RNA accounted for the vast majority of the RNAs isolated from bacteria, and thus ribosomal quantification should serve as an appropriate proxy for bacterial quantification. Using 16S RNA concentration as a proxy for total bacterial RNA concentration may not be appropriate under all circumstances. Ribosome number per cell may vary under certain growth conditions. However, fundamentally, as rRNA comprises the majority of total RNA, total RNA standardization for RT-qPCR is always weighted towards rRNA content. Regardless, for the conditions we have tested, we were able to show that 16S rRNA served as a sufficient proxy for measuring quantities of bacteria, in both infection of naïve plants and under Flg22-pretreated, PTI-preinduced conditions. This allowed us to effectively standardize DC3000 RNA concentrations for downstream analysis. Without correct standardization between samples, the resultant data from RG validation or relative expression measurements may be misleading. We went to great lengths to carefully pair the CFU and 16S rRNA measurements for both methods from the same inoculated leaves, and pairing the data appropriately. We found a linear positive correlation of CFU to 16S rRNA, indicating that bacterial concentration can be calculated based on the quantity of 16S rRNA in a sample.

Using RefFinder, we were able to validate four RGs *(oprF, recA, leuD*, and *rpoD)* for use in this experiment. Our panel of RGs cover normal and high Cq ranges (corresponding to medium and low-expressed transcripts), refer to different cellular functions, and were demonstrated to be stable within the conditions tested in this experiment. However, it is entirely possible that they will not be stable under other experimental conditions or in other hosts. This panel of RGs has been validated for use *in vitro* with King’s B media and for use in Arabidopsis over short time scales of 24 h or less, with and without Flg22 induction of PTI. The use of these RGs under other experimental conditions without validation of stability is not recommended without additional validation. Our analysis indicates that any combination of these four RGs can be used to normalize against a GOI, which affords us some selection in future experiments for GOIs of different ranges of expression or metabolic pathways.

By our calculations, there is often a 10% over-estimation of PCR efficiency when using the serial dilution method as opposed to that derived by LRP. Technical bulletins and the MIQE guidelines (Bustin, Benes et al. 2009) accept up to 110% efficiency, attributing this to user error. Livak’s original 2^-ddCt^ method to calculate relative expression (Livak and Schmittgen 2001) assumed that PCR efficiency was perfect, or at least identical between amplicons. Users of this method gain rough approximates of relative expression, but the equation posed by Pfaffl improved measurements of NRQ because it allows the user to incorporate efficiencies other than 100% (Pfaffl 2001) which allows users to compare relative expression across multiple GOIs. One challenge with the Pfaffl equation was that it could normalize only against a single reference gene, which either requires the user to match their GOIs with inappropriate but historically used RGs, or to seek and validate a new RG for each experiment. The modified Pfaffl equation incorporates multiple RGs, which can satisfy multiple conditions and maintain experimental consistency over time (Hellemans, Mortier et al. 2007). Many software packages exist to assist users in statistical analysis of qPCR, but few can accommodate the entire qPCR data workflow with sufficient statistical rigor, yet remain user-friendly and/or freely available (Pabinger, Rödiger et al. 2014). Our approach utilizes commonly used and free software.

Most users estimate the efficiency by the serial dilution of samples; this is both prone to error and is reagent consuming. Inputting raw amplification values into LRP allows the user to accurately and immediately derive PCR efficiency *in situ* for each sample, which contributes to more precise and conservative measurements of NRQ. This particularly benefits quantitative transcriptomic studies in host/pathogen model systems where the pathogen transcripts are the target of interest, as they are often in low abundance relative to the host’s, and therefore more subject to miscalculation.

As a test of our RT-qPCR method, we examined the NRQ of the virulence-marker genes *hrpA* and *cfl*, as well as *fliC*, a primary PAMP that induces PTI in Arabidopsis. The *hrpA* gene encodes the pilin for the T3SS extracellular appendage used for the delivery of T3SS protein effectors into plant cells (Wei, Plovanich-Jones et al. 2000, MacLean and Studholme 2010). At 5 hpi, *hrpA* expression increases five-fold relative to *in vitro* in infections of both infected and immune-induced plants, suggesting that bacteria are expressing their T3SS and effectors under both treatment conditions. By 24 hpi, NRQs of *hrpA* are reduced, but significantly less so in PTI-induced plants. We speculate that by this time, infection is firmly established in naïve plants, the T3SS is functionaly assembled and effectors have been translocated and T3SS gene expression is no longer necessary. In immune-induced plants, effector delivery is blocked, and the T3SS pilus structural gene is strongly expressed. Together these data suggest that the cues that stimulate down-regulation of T3SS pilus production over the course of an infection are not present or detected in the PTI stimulated plant environment.

The *cfl* gene codes for coronafacic acid ligase, a key enzyme involved in the production of the phytotoxin coronatine, which is a structural mimic of jasmonyl isoleucine and a potent activator of jasmonic acid signaling (Katsir, Schilmiller et al. 2008, Zhang, Yao et al. 2015). Coronatine is associated with the reopening of closed stomata to aid bacterial invasion of leaf tissue (Mittal and Davis 1995, Zheng, Spivey et al. 2012). Five hours into infection, we observed that compared to the inoculum base line, NRQ increased roughly three-fold for *cfl* in bacteria during infection. However, in Flg22-pretreated plants, *cfl* decreased by three-fold on average. By 24 hpi, quantities of cfl *in planta* under both treatments returned to the baseline expression seen the KB-media-raised inoculum, indicating a temporal limit to coronatine synthesis gene expression in this plant/pathogen system.

In contrast, NRQ of *fliC* (flagellin) was repressed by 5 hours and more strongly repressed by 24 hours during infection. However, during PTI exposure, *fliC* was only observed as repressed at 24 hours and the repression was weaker than what was observed during infection. Flagellin is a potent P/MAMP recognized by many plants so it may be evolutionarily advantageous to reduce flagellin expression during plant host interactions to reduce host PTI signaling (Zipfel, Robatzek et al. 2004, Boller and Felix 2009). This is supported by observations of the active repression of flagella seen in *P. syringae* pv. *maculicola* under T3SS inducing conditions (Schreiber and Desveaux 2011). The repression of flagellin may also represent an adaptation by *P. syringae* to prevent mis-targeting of effector proteins for secretion through the flagellar pathway and vice versa (Wei, Chakravarthy et al. 2013).

In conclusion, we have validated a 16S rRNA quantification as an effective approach for standardizing the amount of DC3000 RNA collected from inoculated tissue samples from diverse treatments and time points. We also validated a set of four stable DC3000 reference genes for RT-qPCR expression normalization of DC3000 transcripts during PTI exposure and infection of Arabidopsis. A detailed protocol for this approach is provided in the supplementary text. We have used our approach to demonstrate that PTI exposure alters the expression of key DC3000 virulence-associated pathways. It is our hope that this approach will find utility among researchers seeking to discern the mechanisms of PTI action and support efforts at global transcriptomics analysis of this important model plant pathogen.

## Materials and Methods

### Culture Preparation

Inoculum was prepared by streaking fresh *P. syringae* pv. tomato DC3000 colonies onto a King’s B (KB) agar with 60μg/ml Rifampicin (King E. O. 1954), incubating in the dark for one day at room temperature, smearing the culture in 0.4 ml liquid media to create a lawn, followed by similar incubation. A small patch of lawn was suspended into 0.25 mM MgCl2, and the OD600 brought to 0.9, equivalent to 1x10^9^ CFU/ml, using a Biospectrometer (Eppendorf, Hamburg, Germany). This suspension was further diluted in 0.25mM MgCl_2_ to be used as inoculum. Actual concentration of inoculum was obtained by serial dilution and plating onto KB media with rifampicin. Plates were incubated at room temperature for 24 h, and then moved to 4 °C for three days before counting visible colonies to calculate CFU/ml.

### Infected plant tissue preparation

Fully expanded leaves of four five-week old *Arabidopsis thaliana* Col-0 plants (23C 14/10 L/D, low intensity lighting) were infiltrated using a blunt syringae with one of five decreasing inoculum concentrations, from approximately 10^9^ to 10^6^ CFU/ml in 0.25 mM MgCl2. Using biopsy punches (Integra Life Sciences, Plainsboro, USA), four 0.125 cm^2^ discs were excised from both sides of the midveins of four leaves from each plant, for a total of thirty-two punches per plant. Sixteen punches from the left sides of leaves were pooled and weighed together before grinding in 400 μl 0.25 mM MgCl_2_, then serial-diluted and plated on KB+Rf to obtain CFU/mg leaf tissue. The remaining sixteen punches from the right sides of leaves were frozen in liquid nitrogen for RNA extraction. Each plant was considered a biological replicate (n=4).

For collecting samples from PTI-induced leaves, Two four-to five-week old *Arabidopsis thaliana* Col0 plants were pre-treated by leaf infiltration with 1 μM Flg22, in 0.1% DMSO, 0.25 mM MgCl_2_ 20 h prior to infection. An equal number of plants were mock-treated with an equal volume of 0.1% DMSO, 0.25 mM MgCl_2_. Individual leaves were syringe-infiltrated with approximately 2x10^8^ CFU/ml *P. syringae* pv. *tomato* DC3000 in 0.25 mM MgCl_2_. Two whole leaves from two plants per treatment were harvested at 5 and 24 h post-infection. A single 0.125 cm^2^ disc was excised from each leaf using biopsy punches (Integra Life Sciences, Plainsboro, USA); the four punches were pooled and weighed together before grinding in 100 μl 0.25 mM MgCl_2_, then serial-diluted and plated on KB media with rifampicin to obtain CFU/mg leaf tissue. Plates were incubated as above. The remaining portion of the four leaves were also weighed, divided to two homogenizer tubes, and flash-frozen in liquid nitrogen in preparation for RNA extraction. This experiment was repeated twice with three biological replicates each.

### RNA extraction and Reverse-Transcription

Whole leaves and punched leaf discs were homogenized under freezing conditions in a GenoGrinder (SPEX SamplePrep, Metuchen, USA), and RNA was extracted using Trizol Reagent (Thermo Fisher Scientific, Waltham, USA) followed by the Direct-Zol Miniprep or Miniprep Plus kits (Zymo Research, Irvine USA), which included an on-column DNAse I treatment (30 U/column for 15 m). RNA was quantified using the Biospectrometer and μCuvette G1.0 (Eppendorf, Hamburg, Germany), and the two paired RNA extractions were pooled together. The resulting total RNA extracted was therefore a mix of *in planta* bacterial and plant RNAs. For a media-based *in vitro* control, 1 ml of the inoculum source was pelleted and RNA was extracted in an identical manner.

For each sample, 4-8 μg of RNA was reverse-transcribed to cDNA using the qScript cDNA Supermix (Quantabio, Beverly, USA) using 1 μg RNA per 20 μl reaction as template, the maximum quantity recommended by the kit. A small amount of nonreverse-transcribed RNA was similarly diluted to 50 ng/μl, and retained to test for genomic DNA carryover. RT reactions were carried out in the FlexCycleR^2^ (Analytik Jena AG, Jena, Germany), following Quanta’s recommended thermal cycling protocol (5 m at 25 °C, 30 m at 42 °C, 5 m at 85 °C, hold at 4 °C). After reverse transcription, multiple reactions for individual samples were pooled, aliquoted and stored at -80 °C until use. We found that repeat freeze-thaws of pooled, non-aliquoted cDNA resulted in serious reductions in cDNA concentrations as determined by RT-qPCR. Pre-aliquoting neat cDNA in volumes specific to individual tests ensured equal treatment of transcripts across multi-plate experiments. Dilutions of cDNA to be used in RT-qPCR were prepared in sterile water, on ice or cool blocks, and held at 4 °C for up to 24 h.

## RT-qPCR

### Selection of target genes and primer design

Custom RT-qPCR primers were designed to amplify approximately 100 bp fragments of their target genes based on Genbank accessions for *P. syringae* pv. tomato DC3000 (NCBI Reference Sequence: NC_004578.1), with specifications for a melting temperature of 60 °C. For bacterial RNA quantification, a novel 16S primer set was designed to bind in 16S variable regions 1 and 2 for high *P. syringae* specificity. In Pseudomonads, the 16S variable region 1 has been shown to be hypervariable which aided in the design of specific primers (Bodilis, Nsigue-Meilo et al. 2012). A synthetic dsDNA gBlock (Integrated DNA Technologies, Coralville, USA) was used as a standard for quantification of *P. syringae-specific* 16S rRNA. The gBlock included the entire amplicon, with small flanks on either side, for a total length of 125 bp (Table 1). The gBlock was dissolved in 1X TE to 2.5 ng/μl, then further diluted in sterile water to 1 ng/μl and beyond in a 5-fold serial dilution.

To target reference gene candidates, we used previously-published qPCR primers for *hemD* (Freeman, Chen et al. 2013) and *rpoD* (González-Lamothe, El Oirdi et al. 2012), and designed primers to target *gyrA, gap-1* and *recA*. The remaining four RG candidates, *cynT, lcs-1, leuD* and *oprF* were observed as highly expressed and stable in Flg22-pretreated and naïve infected plant tissues from preliminary RNAseq analysis (manuscript in preparation). To target GOIs, we designed primers for *hrpA* and *fliC*, and used previously published primers targeting *cfl (Worley, Russell et al. 2013)* to amplify under similar conditions (Table 1). Primers were ordered from IDT (Integrated DNA Technologies, Coralville, USA), dissolved in 1X TE to 100 μM, and then diluted to 10 μM in sterile water for use in PCR. All working stocks of primers and the gBlock were pre-aliquoted for individual use and stored at −20 °C.

## RT-qPCR pre-tests

Conditions of RT-qPCR were kept identical throughout all runs. Amplification of cDNA was performed in 10 μl reactions using Veriquest SYBR Green qPCR Master Mix (Thermo Fisher Scientific, Waltham, USA), 0.25 uM primers, and 2 μl cDNA at varying concentrations, depending on treatment and/or target. Master mixes and primers were pre-aliquoted for single use, and stored at -20 °C. New, frozen aliquots of sample cDNA were diluted fresh as needed in sterile water for all reactions that could be run in a single day, and were never re-used. PCR cocktails and plates were prepared on ice or cool blocks, and stored in the dark until ready to run, for no more than 12 h. All PCR reactions were run in triplicate wells. We followed the default thermal cycling protocol in the StepOne software v2.3 (Thermo Fisher Scientific, Waltham, USA) with real-time capture of SYBR Green and ROX (passive reference) fluorescence: 10 m at 90 °C, followed by 40 cycles of 95 °C for 15 s and 60 °C for 1 m, with camera capture at the end of each cycle. A melt curve was generated after the 40^th^ cycle using these parameters: 95 °C for 15 s, 60 °C for 1 m, then a slow ramp (0.3 °C/s) to 95 °C, with camera capture. All runs were conducted on the Step One Plus Real-time PCR System (Thermo Fisher Scientific, Waltham, USA).

To estimate the amplification efficiency (E_est_) of the newly designed 16S rRNA primers, a standard curve was generated in Excel 2016 (Microsoft, Redmond, WA) using 1:5 serial dilutions of *in vitro* bacterial cDNA amplified with *P. syringae-specific* 16S rRNA primers. It was found that PCR reactions containing 0.016 – 5.12x10^-6^ ng pure bacterial cDNA could yield an optimal range of quantification cycles (Cq) from 13.6 – 24.9, and the slope of the curve generated by plotting the log of ng cDNA against the resulting C_q_ would estimate a good efficiency in pure bacteria (105.08%, R^2^ = 0.9972), using the formula

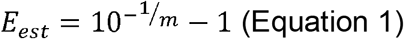

(Supplementary data S1, “16S Standard Curve”).

To test the amplification efficiency of the 16S primers in mixed plant and bacterial cDNA, and to determine relative concentrations of bacterial cDNA derived from mixed total RNAs, we infected plants as described above with decreasing concentrations of inoculum (approx. 1x10^9^, 3x10^8^, 1x10^8^, 3x10^7^, 1x10^7^, and 3x10^6^ CFU/ml) for 2 h, and performed RNA extractions and RT synthesis as described above. cDNA was serially diluted 1:5 from 2 ng/μl, and three dilutions from 0.4 – 0.016 ng per reaction were tested with the 16S primers. Cqs were plotted against the log10 of ng cDNA, and the Eest was calculated from the slope of the curves. All six samples showed Eest within somewhat acceptable ranges (94 – 114%), with good R^2^ values (0.9911 – 1) for correlation coefficients (Supplementary data S1, “Eest of 16S in var. inoculum”).

To verify that all endogenous gene primer pairs used in the study showed 90-110% efficiency, pure bacterial cDNA was serially diluted 1:5 from 10 and 2ng/μl to create a range of concentrations from 20.0 – 0.0032ng per reaction. A series of these dilutions were amplified with each of nine candidate RG primer sets (Supplementary data S1, “RG pre-tests”).

### Sample quantification and reference gene validation

For RT-qPCR, a new 8-point standard curve was generated using 1/5^th^ serial dilutions the 16S gBlock amplified with the *P. syringae-specific* 16S rRNA primers, ranging from 0.016 – 2.05x10^-7^ng cDNA per reaction. *In planta* and *in vitro* samples were diluted to a concentration appropriate to detect within a normal Cq range (approximately Cq = 20 – 24), as found by preliminary experiments. The bacterial content of all samples was quantified by plotting against the 16S standard curve. Concerns regarding carryover of genomic DNA into the RT-qPCR reaction were eliminated by testing equivalent amounts of non-reverse-transcribed RNA. Bacterial cDNA quantities detected in the PCR were scaled up to the amount of total RNA converted to cDNA in the first-strand reverse-transcription synthesis. Total cDNA was further scaled up to quantities of extracted total RNA, and the weight of plant tissue used was factored. These quantities in ng 16S rRNA per mg leaf weight were plotted against recovered CFU per mg leaf weight to show equivalence in detection by plating and PCR (Figure 1A and 1B).

To investigate the variation of expression of candidate reference genes across samples generated under different conditions, all samples were normalized to 0.6 pg bacterial rRNA per reaction, a concentration slightly lower than the least-concentrated sample. The normalized samples were tested against all nine RG primer pairs, and a 1:1250 dilution was tested against the 16S primer pairs as well to confirm normalization. Data were trimmed to remove outliers within the StepOne software: individual reactions that contributed to an average C_q_ with a standard deviation (SD) higher than 0.3 were removed. Individual wells that showed aberrant amplification curves, especially those in corner wells or with very high starting concentrations of template, were also removed, even if their SD was below 0.3, because downstream analysis may derive incorrect baseline values from them. To determine quantification cycle (Cq) and calculate empiric PCR efficiency (E_emp_), raw amplification data (Rn values) were input to LinRegPCR (LRP), version January2016, available from the Heart Failure Research Center. Baselines were determined according to default settings, and the Window-of-Linearity (WOL) was set to establish a baseline via linear regression for each amplicon. If samples showed baseline errors, the errors were ignored and samples were allowed to remain in the dataset, but would not contribute to the mean efficiency as derived by the LRP software (E_emp_). The mean C_q_ values of triplicate wells for each RG and sample were input to RefFinder, available from the Cotton EST Database. This software ranks RGs across multiple algorithms for stability (Figure 2). From these tests, we were able to select our four top RGs, *oprF, recA, leuD, rpoD*. We chose to use *oprF* and *recA* in subsequent PCRs because they demonstrated normal *(oprF)* and late (recA) Cq ranges, representing mid- and low-expressed transcripts, respectively.

### Genes of Interest

We tested normalized cDNA from five treatments: *in vitro* bacteria, mock-treated 5 h infecting bacteria, Flg22-pretreated 5 h infecting bacteria, mock-treated 24 h infecting bacteria and Flg22-pretreated 24 h infecting bacteria). We used three biological replicates each from duplicated experiments. One sample in the Flg22, 5 h set was discarded from analysis due to consistently aberrant results, most likely due to incomplete Flg22 pre-treatment. These samples were tested for relative expression of three GOIs: coronafacic acid ligase *(cfl)*, flagellin *(fliC)* and the Type III secretion system pilin protein *(hrpA)*. Samples were grouped by biological replicate set, and tested against the three GOIs, the two RGs, and 16S rRNA within the same plate. Pairing GOIs with RGs and treatment samples to controls should be prioritized over more convenient groupings within a single plate, as variation between runs can complicate all calculations. This became evident when results of identical samples in different runs showed wide swings in Cq values.

### Measurement and Analysis of Relative Expression

Individual data were trimmed in StepOne software to eliminate outliers, and the raw amplification data from these trimmed samples were input to LRP. Using the baseline and WOL settings described above, sample Cq, and mean Eemp were determined. The normalized relative quantity (NRQ) of a GOI was calculated using Eemp and Cqs from LRP, via a modified Pfaffl equation (Pfaffl 2001, Hellemans, Mortier et al. 2007) to include the geometric mean of both RGs in the denominator:

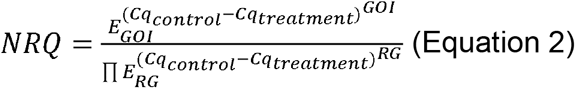

Cqs of three technical replicates were averaged, and the NRQ of five-to-six biological replicates was calculated individually, so that GOI and RG values could be paired appropriately for data collected within runs. For each GOI, we performed two-tailed t-tests assuming equal variances in MATLAB and Statistics Toolbox Release 2012b (The Mathworks, Inc., Natick, USA) to test the null hypothesis that the means of NRQ values between treatments were equal. The null hypotheses of t-tests were rejected at the 5% significance level (p<0.05) (Figure 3).

## Acknowledgements

We gratefully acknowledge the assistance of Dr. Ian Major with statistical guidance, Dr. Marin Brewer for use of the Genogrinder and Drs. Ian Major and Bryan Swingle for helpful discussion and comments regarding the preparation of this manuscript.

